# Baculoviral COVID-19 Delta DNA vaccine cross-protects against SARS-CoV2 variants in K18-ACE2 transgenic mice

**DOI:** 10.1101/2022.10.07.511252

**Authors:** Yuyeon Jang, Hansam Cho, Jungmin Chun, Kihoon Park, Aleksandra Nowakowska, Jinha Kim, Hyeondong Lee, Chanyeong Lee, Yejo Han, Hee-Jung Lee, Ha-Youn Shin, Young Bong Kim

**Affiliations:** Department of Bio-industrial Technologies, Konkuk University, 120 Neungdong-ro, Gwangjin-gu, Seoul, Republic of Korea; KR BioTech, 120 Neungdong-ro, Gwangjin-gu, Seoul, Republic of Korea; Department of Bio-medical Science and Engineering, Konkuk University

**Author notes:** Corresponding Author: Prof. Young Bong Kim.

**Keywords:** SARS-CoV-2, SARS-CoV-2 vaccine, Baculoviral DNA vaccine, SARS-CoV-2 Variants, AcHERV-COVID19 Delta vaccine, Cross protection

## Abstract

After severe acute respiratory syndrome coronavirus-2 (SARS-CoV2) made the world tremble with a global pandemic, SARS-CoV2 vaccines were developed. However, due to the coronavirus’s intrinsic nature, new variants emerged, such as Delta and Omicron, refractory to the vaccines derived using the original Wuhan strain. We developed an HERV-enveloped recombinant baculoviral DNA vaccine against SARS-CoV2 (AcHERV-COVID19S). A non-replicating recombinant baculovirus that delivers the SARS-CoV2 spike gene showed a protective effect against the homologous challenge in a K18-hACE2 Tg mice model; however, it offered only a 50% survival rate against the SARS-CoV2 Delta variant. Therefore, we further developed the AcHERV-COVID19 Delta vaccine (AcHERV-COVID19D). Cross-protection experiments revealed that mice vaccinated with the AcHERV-COVID19D showed 100% survival upon challenge with Delta and Omicron variants and 71.4% survival against prototype SARS-CoV2. These results support the potential of the viral vector vaccine, AcHERV-COVID19D, in preventing the spread of coronavirus variants such as Omicron and SARS-CoV2 variants.

**Author Summary:** After the SARS-CoV2 pandemic, it is known that the existing vaccine has diminished efficacy against the emerging variants. We developed a baculoviral COVID19 DNA vaccine for the Delta variant (AcHERV-COVIS19D). Compared to AcHERV-COVID19S, designed to protect from the prototype of SARS-CoV2, AcHERV-COVID19D elicited higher humoral and cellular immunity and showed perfect protection against SARS-CoV2 delta strain and Omicron challenge. The broad and robust cellular immunity of the AcHERV-COVID19D vaccine appears to have played a significant role in the cross-protection of the Omicron variant. Our AcHERV-COVID19D can be a potential vaccine against emerging SARS-CoV2 variants.

## Introduction

Severe acute respiratory syndrome coronavirus 2 (SARS-CoV2) was declared a pandemic by the World Health Organization (WHO) on 11 March 2020. SARS-CoV2 vaccines were rapidly developed following the global emergence of this pandemic, and the vaccine benefit has been shown to significantly lower mortality rates (1–3). However, the emergence of SARS-CoV2 variants of concern (VOC) reduced potency of existing prototype vaccines are driving the development of vaccines that could provide protection against VOC (4–8).

We previously developed a SARS-CoV2 prototype vaccine using a baculoviral vector system and evaluated the efficacy of the developed AcHERV-COVID19S prototype vaccine in Syrian golden hamsters (9). In this study, we confirmed the efficacy of the AcHERV-COVID19S vaccine in K18-hACE2 transgenic mice. Additionally, we developed the AcHERV-COVID19D vaccine that delivers the spike gene of the SARS-CoV2 Delta variant. The primary focus of this study was to assess the efficacy of the AcHERV-COVID19D against the SARS-CoV2 Delta variant. Given the rising prevalence of the Omicron variant, we further assessed the cross-protection ability of the AcHERV-COVID19D vaccine against this variant in K18-hACE2 transgenic mice.

Developing customized vaccines against new variants has a limitation in that it cannot keep up with new VOCs due to the need for long-term vaccine development. Therefore, we also performed a cross-protection evaluation against the SARS-CoV2 prototype, Delta, and Omicron variants to study the potential of AcHERV-COVID19D as a vaccine against emerging VOCs.

## Results

### Construction and immunogenicity of AcHERV-COVID19 vaccines

A baculovirus-based COVID19 DNA vaccine, the AcHERV-COVID19 prototype vaccine (termed AcHERV-COVID19S) was constructed to deliver the spike gene of the SARS-CoV2 prototype (Wuhan). To develop the recombinant baculovirus containing the SARS-CoV2 Delta variant, the RBD-to-S1 sequence of the AcHERV-COVID19 prototype was replaced with that from the Delta variant termed AcHERV-COVID19D. To enhance the immunogenicity of the spike protein, we substituted two prolines at residues K986 and V987 and removed the polybasic cleavage site by R685N (RRAR to RRAN), as suggested in a previous report (10). A schematic diagram of the recombinant baculoviruses encoding SARS-CoV2 is shown in Figure 1A. To characterise S antigen expression, Vero E6 cells infected with AcHERV-COVID19S or AcHERV-COVID19D were evaluated by immunofluorescence assay and Western blotting using spike-specific antibodies (Fig. 1B, C).

**Fig. 1.**
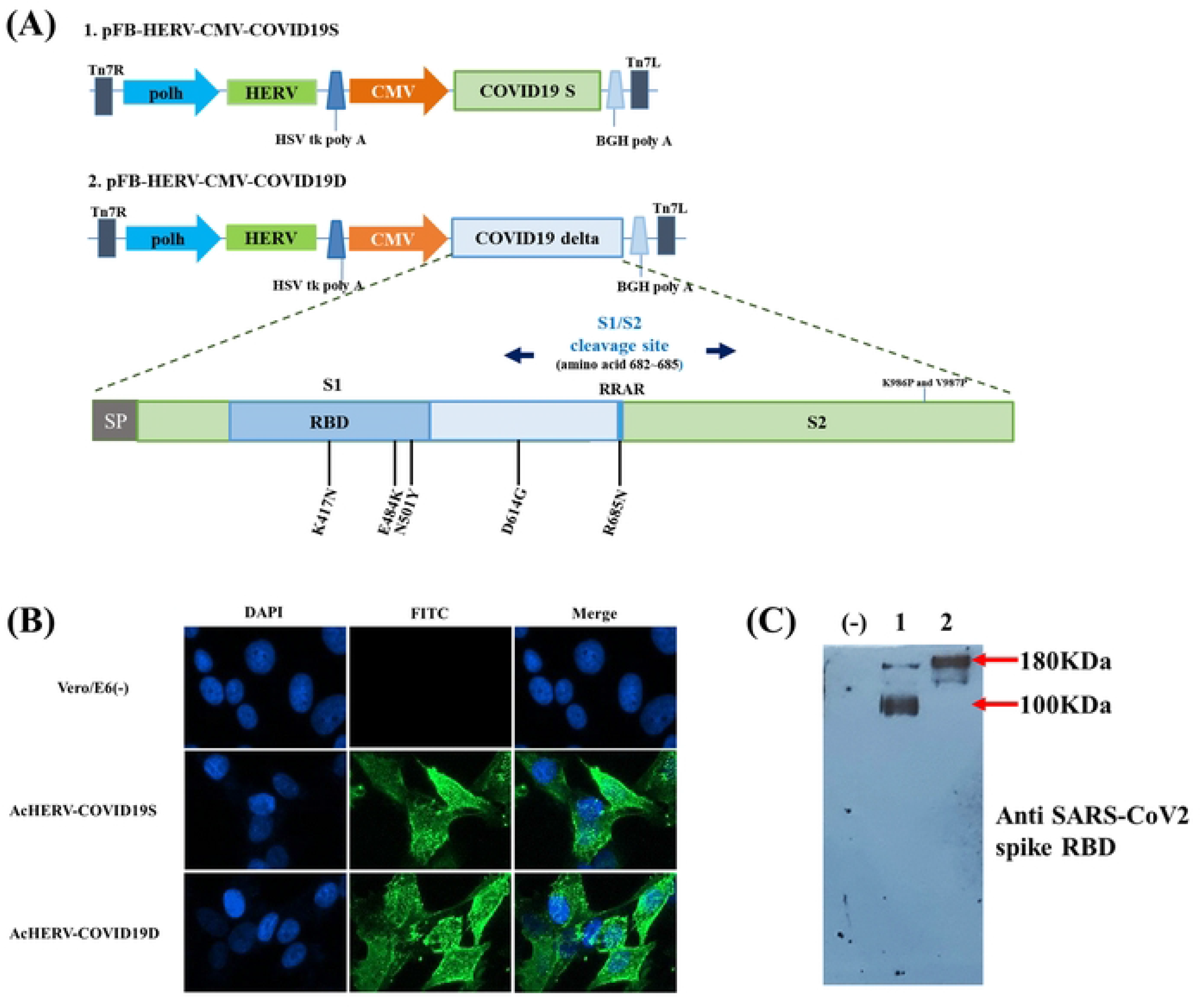
Construction and immunogenicity of AcHERV-COVID19 vaccines. (A) Schematic diagrams of the COVID19 recombinant baculoviruses. Expression of SARS-CoV2 S protein was detected by immunofluorescence assay (B) and Western blotting (C) in Vero E6 cells infected with AcHERV-COVID19S and AcHERV-COVID19D. (-): uninfected cells; Lane 1: AcHERV-COVID19S; Lane 2: AcHERV- COVID19D.

### AcHERV-COVID19S protects against SARS-CoV2 challenge in K18-hACE2 mice

To evaluate the immunogenicity of the AcHERV-COVID19S (prototype), Six-week-old female K18-ACE2 mice were immunized with AcHERV-COVID19S twice intramuscularly (Fig. 2A, S1 Table). Two weeks after boosting vaccination, all vaccinated mice showed induction of SARS-CoV2-specific total IgG levels (Fig. 2 B). But the level of neutralising antibodies was not as high as seen in the serum of an individual who had received commercial vaccines (positive control) (Fig. 2C).

**Fig. 2.**
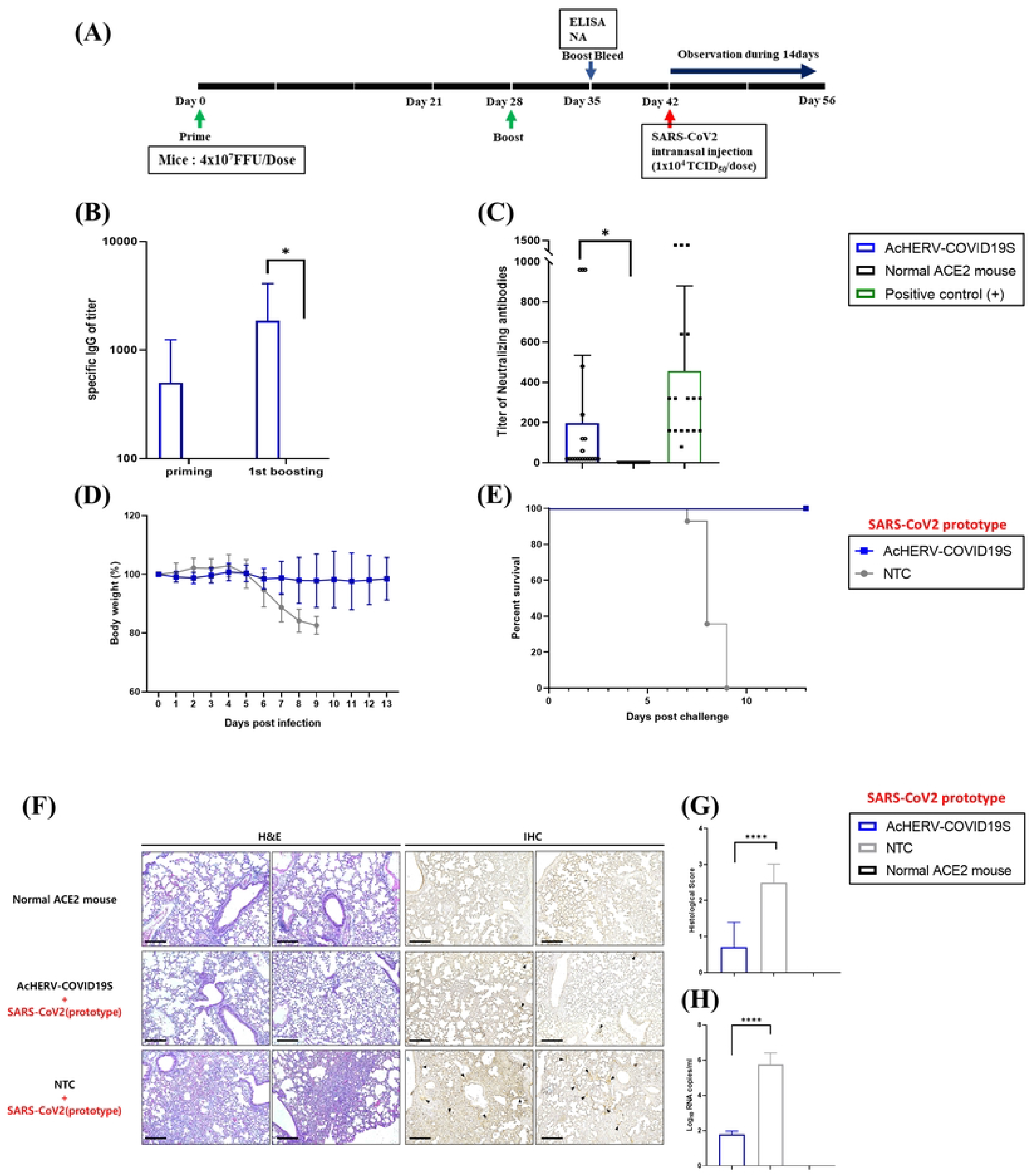
Immunogenicity of AcHERV-COVID19S in K18-ACE2 mice and SARS-CoV2 prototype challenge experiment. (A) Immunization schedules. (B) Detection of IgG antibody responses by ELISA. (C) Neutralizing antibodies against the SARS-CoV2 prototype. (D) Body weight. (E) Mouse survival rate. (F) H&E staining and Immunohistochemistry. Scale bar = 100 μm. (G) Histological evaluation. It was based on the percentage of inflammation area from each animal using the following scoring system in infected mice (i.e., 0 = absent, 1 = minimal, 2 = moderate, or 3 = severe). (H) Viral lung titres in lungs on 7 days after challenge. *P*-values were generated using a ANOVA followed by the Tukey–Kramer post-analysis test. ****: P < 0.0001.

To test the prophylactic effect of the AcHERV-COVID19S, mice were challenged with a lethal dose of SARS-CoV2 (prototype). After SARS-CoV2 infection, the mice from the non-treatment control (NTC) group lost more than 17.4% of their initial body weight, and all died on the 9th day. The AcHERV-COVID19S vaccine group showed mild weight loss and symptoms but recovered rapidly over 13 days and exhibited 100% survival with good health (Fig. 2D-E).

The results of histopathological analysis of mouse lung tissues are shown in Figure 2F-G. Seven days after infection, lung tissues of the AcHERV-COVID19S vaccine group showed a significantly milder pathology upon SARS-CoV2 infection. Unlike the vaccine group, NTC mice showed severe lesions, perivascular and alveolar infiltration, and severe inflammation compared to the lungs of normal mice. On day 13 post-challenge, viral titres were measured from the lung tissues of all surviving mice using qRT-PCR. The virus was weakly detected in mice at this point but was below the lower limit of detection in some cases (Fig. 2H). Thus, although AcHERV-COVID19S vaccinated mice did not show complete sterile protection, all such mice recovered and survived.

### Cell-mediated immunity plays an important role in the recovery after SARS-CoV2 infection

To measure the cellular immunity-inducing effects of the vaccine, an additional experiment was performed using C57BL/6 mice. An ELISPOT assay revealed that the level of IFN-γ-secreting splenocytes was significantly higher in AcHERV-COVID19S vaccinated mice compared to control mice (*: *P*<0.05, S1 Fig A). We also measured the induction of Th1- and Th2-type cytokines by qRT-PCR. TNF-α and IL-2 were used to assess Th1-type levels, while IL-4 was used to assess Th2-type cytokines. We found that the mRNA expression levels of TNF-α, IL-2, and IL-4 were higher in the immunized group than in the NTC group. The vaccinated group showed greater Th1 cell immune responses and maintained higher Th1 cytokine mRNA levels than the NTC group (S1 Fig. B). This finding demonstrated importance of the Th1-skewed immune response in protecting against infection and reducing the clinical severity of subsequent infections (11). Collectively, these data suggest that the ability of AcHERV-COVID19S to induce strong cellular immunity plays an important role in the recovery of vaccinated K18-hACE2 transgenic mice after SARS-CoV2 infection.

### Immunogenicity comparison of AcHERV-COVID19S prototype and AcHERV-COVID19D delta vaccine

To compare the immunogenicity of the AcHERV-COVID19D delta vaccine with that of the AcHERV-COVID19S (prototype vaccine), SARS-CoV2-specific humoral immune responses were evaluated in K18-ACE2 mice. K18-ACE2 mice were immunized twice intramuscular injection into the hind legs with AcHERV-COVID19S or AcHERV-COVID19D at a 4-week interval (Fig. 3A, S2 Table). Compared to the AcHERV-COVID19S, the AcHERV-COVID19D delta vaccine showed significantly improved SARS-CoV2 S-specific IgG titres (Fig. 3B; AcHERV-COVID19S vs. AcHERV-COVID19D *: P < 0.05). Two weeks after the second vaccination, the neutralizing antibody titres of sera from AcHERV-COVID19D and AcHERV-COVID19S-immunised mice were evaluated against SARS-CoV2 prototype and Delta variant viruses. As expected, the sera of AcHERV-COVID19D immunized mice had a high neutralizing antibody titre against the homologous Delta virus, which was higher than that of AcHERV-COVID19S immunized mouse sera. In contrast, the heterologous neutralizing antibody titre of delta vaccine-immunized mice was much lower (by about 4-fold) for the prototype SARS-CoV2 virus (Fig. 3C).

**Fig. 3.**
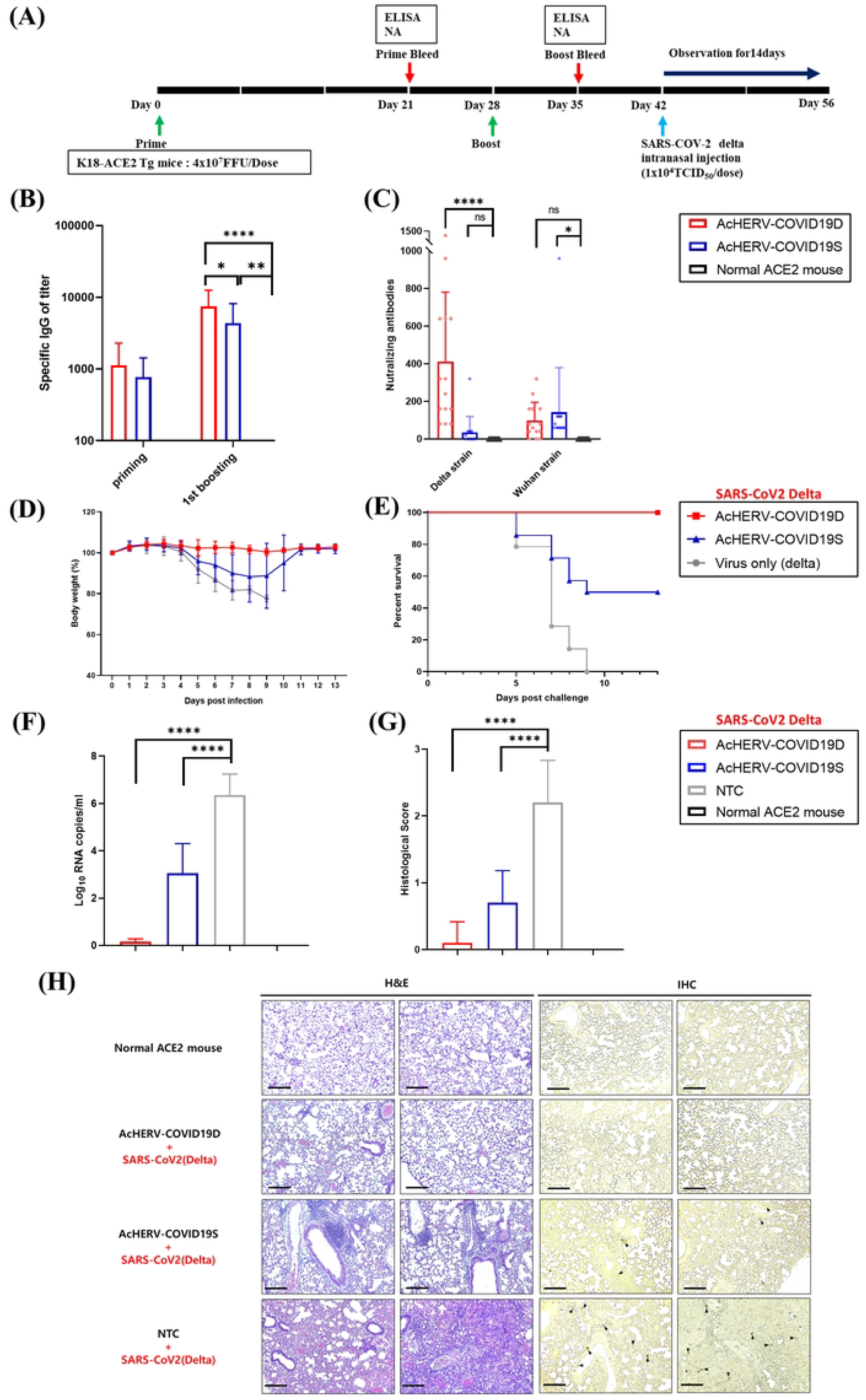
Immunogenicity of AcHERV-COVID19S and AcHERV-COVID19D in K18-ACE2 mice and SARS-CoV2 Delta challenge experiment. (A) Immunization schedules. (B) Detection of IgG antibody responses by ELISA. (C) Neutralizing antibodies against the SARS-CoV2 Delta virus. (D) Body weight. (E) Survival rate. (F) Viral lung titres on 7 days after challenge. (G) Histological evaluation. (H) H&E staining and IHC in lungs. Scale bar = 100 μm. P-values were generated using an unpaired two-tailed t-test or ANOVA followed by the Tukey–Kramer post-analysis test. NS: P > 0.05, *: P < 0.05, **: P < 0.01, ****: P < 0.0001.

### AcHERV-COVID19D can protect against the homologous pathogenic SARS-CoV2 Delta variant

To test the prophylactic effects of the AcHERV-COVID19D, mice immunized with AcHERV-COVID19S or AcHERV-COVID19D were challenged with the SARS-CoV2 Delta variant. After the challenge, the mice of the NTC group lost more than 22.35% of their initial body weight, and all died by day 9. In contrast, the mice from the AcHERV-COVID19D delta group (N=14) appeared healthy, with maintained weight, and all survived SARS-CoV2 Delta infection until the study endpoint (day 13) (Fig. 3D-E). The AcHERV-COVID19S group (N = 14) exhibited weight loss, symptoms, and 50% mortality (7 of 14 mice). This suggests that the AcHERV-COVID19S vaccine did not effectively offer cross-protection against SARS-CoV2 Delta infection.

On day 13, all surviving mice were humanely euthanized, lungs were isolated, and the virus titre was measured (Lungs of NTC were isolated on day 9). Notably, there was no detectable virus in the lungs of mice immunized with AcHERV-COVID19D. In contrast, high virus titres were present in the lungs of mice from the AcHERV-COVID19S and NTC groups (Fig. 3F). Additionally, histopathological analysis of lungs from vaccinated mice showed reduced histopathological symptoms, and histological scores also showed a similar decrease (Fig. 3G-H).

Together, these results indicate that AcHERV-COVID19D vaccine showed perfect protection against the homologous pathogenic SARS-CoV2 Delta variant without the lung inflammation and/or injury associated with SARS-CoV2 infection.

### Cross-protection evaluation of AcHERV-COVID-19D DNA vaccine in K18-ACE2 Tg mice

As new SARS-CoV2 variants emerged, we performed Omicron cross-protection tests using the AcHERV-COVID19D vaccines. For this experiment, replication-incompetent adenoviral vector vaccines were constructed as a control to deliver the spike genes for the SARS-CoV2 prototype or Delta variants. The principle underlying this vaccine is the same as that of the AstraZeneca vaccine, and its characterization is described in S2 Fig.

To examine the cross-reactivity to the AcHERV-COVID19D vaccine, 112 mice were divided into 13 groups, as shown in Table 1, and an experiment was carried out. Groups 2, 7, and 11 (total n=28) were immunized with the AcHERV-COVID19D delta vaccine and challenged with three different SARS-CoV2 viruses. Groups 3 and 8 (each n=7) immunized with adenovirus vectored COVID19S vaccine were challenged with SARS-COV2 Delta and prototype, respectively. Groups 4, 9, and 12 (each n=7) immunized with adenovirus vectored COVID19D vaccine were challenged with SARS-CoV2 Delta and Omicron, respectively.

**Table 1.**
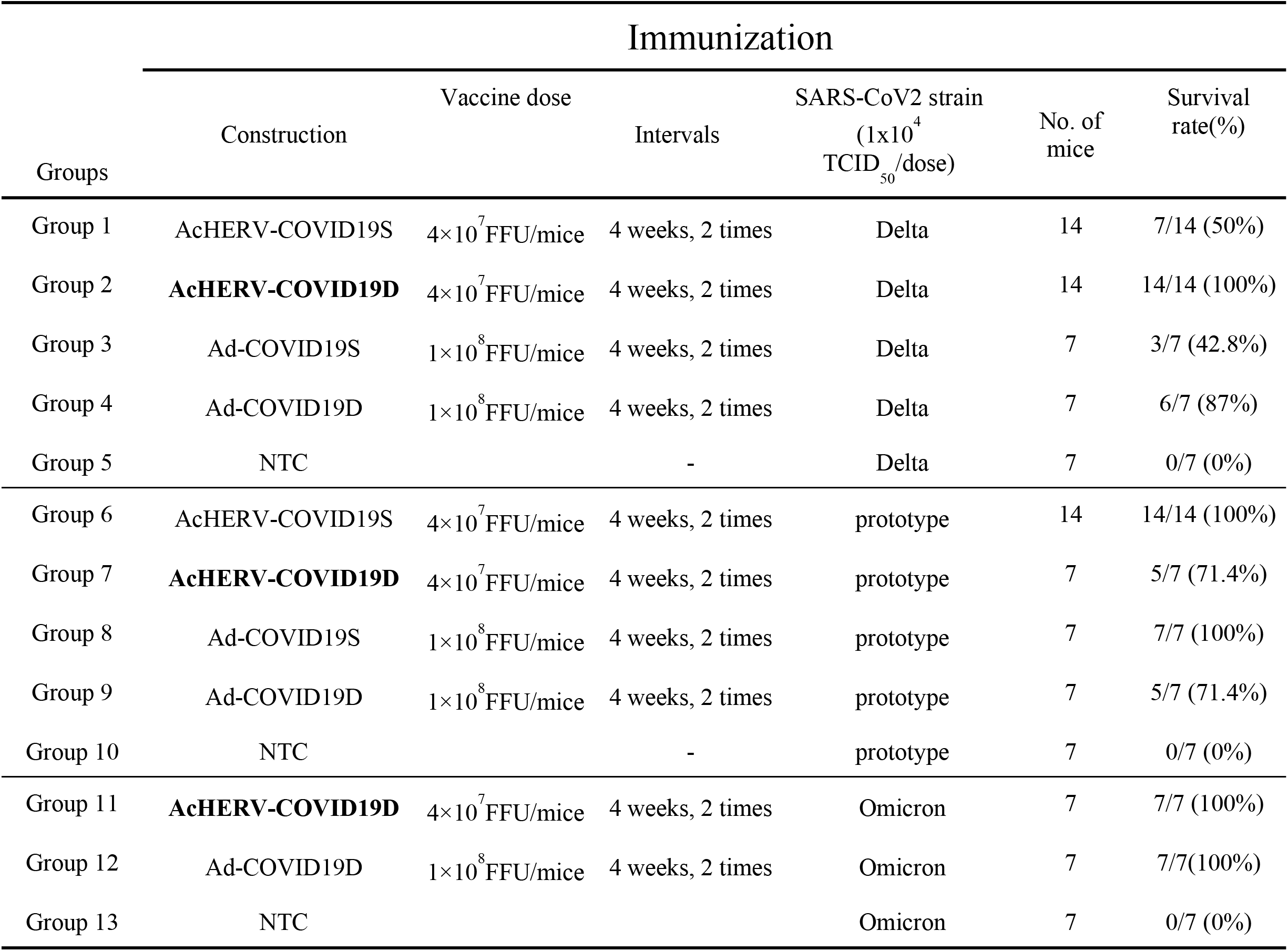
Cross-protection Experiment against various SARS-CoV-2 strains.

Figure 4A shows the neutralizing antibody titres against the prototype, Delta, and Omicron viruses in the sera of 28 mice immunized with the AcHERV-COVID19D (Groups 2, 7, 11). The maximum neutralizing antibody titre of over 1,280 was observed against the Delta virus. In contrast, titres of 640 and 320 were obtained for the prototype and Omicron SARS-CoV2 strains, respectively (Fig. 4A). To measure the cellular immunity of the vaccine against various SARS-CoV2 strains, an additional experiment was performed using the ELISPOT assay. Mice vaccinated with AcHERV-COVID19D showed significantly higher levels of IFN-γ secreted from splenocytes against three SARS-CoV2 strains without a significant difference (Fig. 4B).

**Fig. 4.**
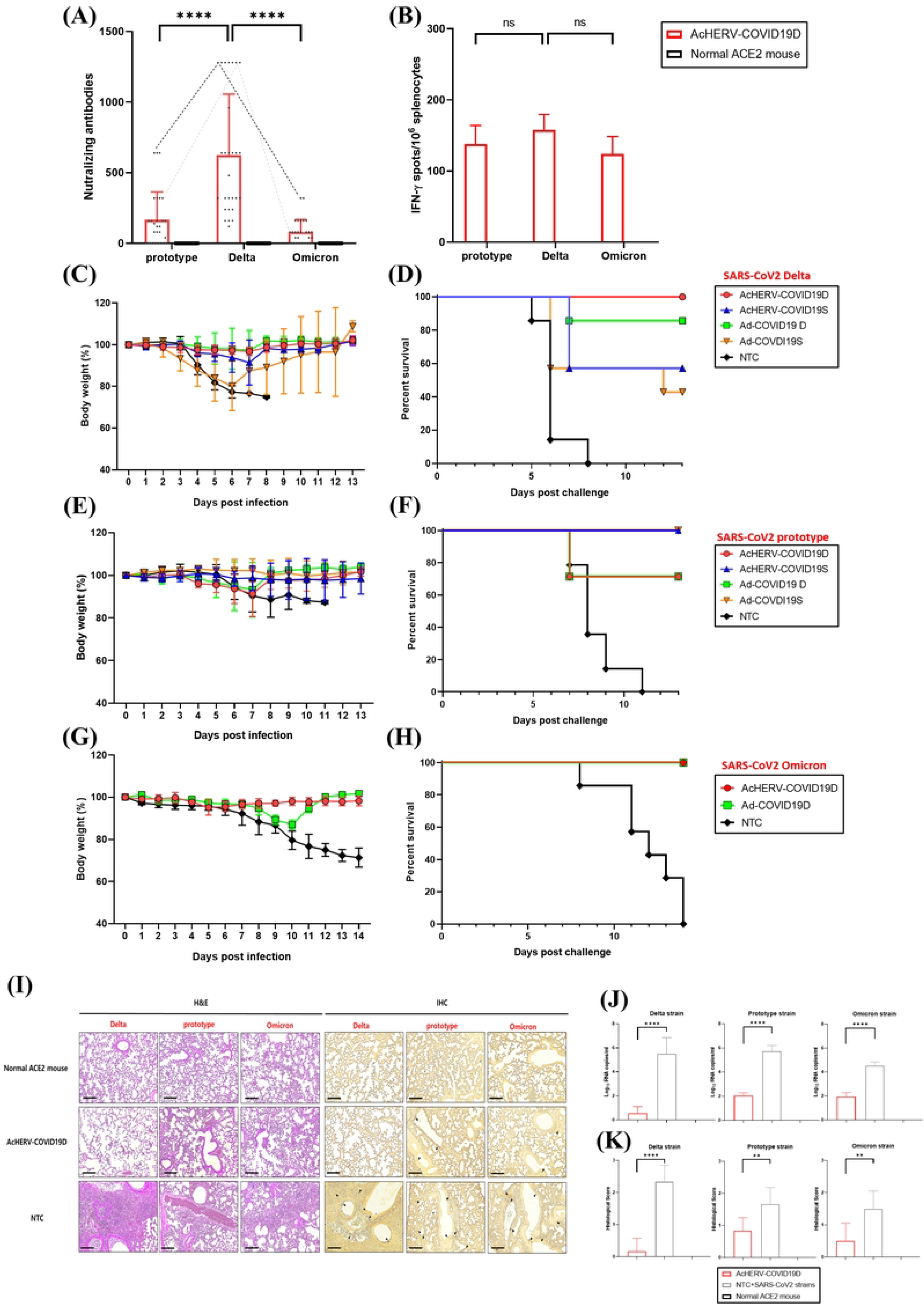
Comparison of immunogenicity and protection against various SARS-CoV2 strains in K18-ACE2 mice. Comparison of various SARS-CoV2 strain-specific neutralizing antibody responses (A) and ELISPOT assay (B) in mice immunized with AcHERV-COVID19D. Body weight (C, E, G) and survival rate (D, F, H) challenged with SARS-CoV2 Delta, prototype and Omicron, respectively. (I) H&E and IHC staining. Histological evaluation was tested in the lungs of the group infected with the SARS-CoV2 Delta, Prototype, or Omicron. Scale bar = 100 μm. (J) Viral lung titres of various strains in vaccinated mice and NTC were measured by qPCR. (K) Histological scores. P-values were generated using a ANOVA followed by the Tukey–Kramer post-analysis test. NS: P > 0.05, **: P < 0.01, ****: P < 0.0001.

Figure 4C–4H shows the body weight and survival rate against the prototype, Delta, and Omicron viruses in mice immunized with the AcHERV-COVID19D. The AcHERV-COVID19D showed 71.4%, 100%, and 100% protection in mice challenged with the prototype, Delta, and Omicron variant, respectively.

Figures 4I–4K show histopathological analysis and viral load for prototype, Delta, and Omicron viruses in mice immunized with AcHERV-COVID19D. The pathological results of lung tissue in the AcHERV-COVID19D vaccinated group were weaker lesions and significantly reduced viral load compared to the NTC group. The body weight, survival rate, histological analysis, and viral load results confirmed that our AcHERV-COVID19D vaccine provided perfect protection against the Delta virus. We also confirmed a high level of protection against the prototype virus. In the case of the Omicron variant, the mice in the NTC group died on day 14, showing heavy viral load and lesions in histological analysis. The AcHERV-COVID19D vaccine group showed a 100% survival rate against the Omicron virus. The histological examination showed a low lesion phenomenon and a significantly low viral load.

## Discussion

We had previously developed MERS-CoV, SARS-CoV2, and Zika virus vaccines using a baculovirus-based DNA vaccine (AcHERV) system (9, 12). Here, we demonstrated that AcHERV-COVID19D could be a potential SARS-CoV2 vaccine against pathogenic Delta and Omicron variants. Polybasic cleavage site-modified AcHERV-COVID19D elicited stronger humoral and cellular immunity than the AcHERV-COVID19S. Although the AcHERV-COVID19S vaccine induced low NA, vaccinated mice showed 100% survival against homologous SARS-CoV2 virus challenge. These findings suggest that the cellular immunity conferred by AcHERV-COVID19S played a vital role in the recovery of mice after SARS-CoV2 infection. This ability to induce cellular immunity can be an advantage of mRNA and viral-vector DNA vaccines (2, 13, 14). However, the non-modified AcHERV-COVID19S vaccine showed a 50% survival rate against highly pathogenic heterologous SARS-CoV2 Delta. This indicates that humoral immunity is also very important against the virulent Delta variant, emphasizing that vaccines should seek to induce both cellular and humoral immunity.

The continuous emergence of VOCs has become the biggest problem in efforts to develop vaccines against SARS-CoV2. Notably, AcHERV-COVID19D conferred 100% protection in mice challenged with the Omicron variant, whose spike protein differs from that of Delta by 34 amino acids (15, 16). We speculate that the cross-protection ability of AcHERV-COVID19D may be related to the virulence of a given virus. The ability of the AcHERV-COVID19D to cross-protect against Omicron is beyond the level expected for the observed neutralizing antibody activity (Fig. 4A), indicating that cellular immunity is involved (Fig. 4B).

Although weight loss was observed, the adenoviral delta vaccine (Ad-COVID19D) group also showed 100% survival following the Omicron challenge. The Ad-COVID19D is replication-incompetent in mice, it expresses all adenoviral genes except for the IE3 gene and thus exerts viral toxicity. In contrast, a baculovirus vector vaccine is non-toxic; most baculoviral gene promoters are not recognized by RNA polymerase II, and thus the virus-derived proteins cannot be expressed (17, 18). Also, because of this silence, there are no challenges associated with vector antibodies. Thus, the AcHERV vaccine system could be implemented for developing a booster vaccine.

In conclusion, AcHERV-COVID19D elicited higher humoral and cellular immunity and showed perfect protection against SARS-CoV2 delta strain and Omicron challenge. The broad and robust cellular immunity of the AcHERV-COVID19D vaccine appears to have played a significant role in the cross-protection of the Omicron variant. Our AcHERV-COVID19D can be a potential vaccine against emerging SARS-CoV2 variants.

## Material & Methods

### Viruses and cells

*Spodoptera frugiperda* 9 (Sf9, Invitrogen, USA) cells were propagated in Sf-900II medium (Invitrogen, USA) supplemented with 1% antibiotic-antimycotic (Invitrogen, USA) at 27°C. VeroE6 (ATCC, USA) cells were cultured in DMEM supplemented with 10% FBS (Gibco, USA) at 37°C.

SARS-CoV2 prototype (BetaCoV/Korea/LCDC03/2020, NCCP No.43326), SARS-CoV2 Delta B.1.617.2 (NCCP No.43390), and SARS-CoV2 Omicron BA.2 (NCCP No.43408) were obtained from the National Culture Collection for Pathogens (Korea Disease Control and Prevention Agency). All experiments were performed in the Biosafety Level 3 facility of Konkuk University (Republic of Korea, KUIBC-2021-01, KIBIC-2022-01).

### Construction of recombinant baculoviruses (AcHERV-COVID19D)

A recombinant baculovirus encoding full-length SARS-CoV2 was previously constructed by inserting the sequence into pFastBac-HERV. To construct AcHERV-COVID19D, the fragment spanning from the RBD sequence to the S1 coding sequence for the SARS-CoV2 Delta S protein was amplified using PCR. Residues K986 and V987 were replaced with prolines to stabilize the pre-fusion form of S. The S1/S2 cleavage site of the spike protein was replaced with asparagine residue (RRAR to RRAN). Finally, part of the RBD-to-S1 sequence of the original COVID19 S was replaced with that of the Delta variant to generate the AcHERV-COVID19D lacking the cleavage site (10). Recombinant baculoviruses were produced using the Bac-to-Bac baculovirus expression system according to the manufacturer’s manual (Invitrogen).The scheme for constructing the recombinant baculoviruses, AcHERV-COVID19S, and AcHERV-COVID19D, is shown in Fig. 1A.

A recombinant adenovirus delivering the full-length SARS-CoV2 S gene was constructed by inserting the relevant sequence into pAdenoX-CMV (S2 Fig 2). Recombinant adenovirus vaccines were produced using the Adeno-X™ Adenoviral System 3 according to the manufacturer’s manual (Takara Korea Biomedical Inc., Republic of Korea).

### Immunofluorescence assay and Western blotting

The expression of SARS-CoV2 S prototype and S Delta variant proteins in mammalian cells was tested by infecting Vero E6 cells with baculovirus at a multiplicity of infection of 30. Three days after infection, immunofluorescence analysis and Western blotting were performed as described previously (9) using a SARS-CoV2 polyclonal antibody (Elabscience, USA).

### Animals

B6.cg-Tg (K18-hACE2) 2Prlmn/J mice were purchased from Jackson Laboratory (USA), and 6-week-old female C57BL/6 mice were purchased from Orient-Bio (Republic of Korea). The mice were supplied food and water ad libitum. The SARS-CoV2 challenge experiment was performed in the Animal Biosafety Level 3 facility. All animal experiments were approved by the Konkuk University Institutional Animal Care and Use Committee (IACUC approval No: KU21160) and were conducted strictly in compliance with the Guide for Care and Use of Laboratory Animals of the National Institute of Health.

### Immunization

Female C57BL/6 and B6.cg-Tg (K18-ACE2) 2Prlmn/J mice were immunized by intramuscular injection into the hind legs with 4 × 10^7^ FFU of AcHERV-COVID19S or AcHERV-COVID19D delta. K18-ACE2 mice were immunized by intramuscular injection into the hind legs with 1 × 10^8^ IFU of recombinant adenovirus (Ad-COVID19S or Ad-COVID19D). Mice were immunized twice at a 4-week interval. Blood samples were collected after mice were anesthetized by intramuscular injection with 40 mg/kg of Zoletil 50 (Virbac Laboratories, France) and 5 mg/kg of Rompun (Bayer Korea, Republic of Korea).

### Enzyme-linked immunosorbent assay

Induction of antibodies specific for SARS-CoV2 S was tested by enzyme-linked immunosorbent assay (ELISA). To detect the SARS-CoV2 S-specific antibody, a 96-well plate was coated with the SARS-CoV2 S RBD protein (produced in the Sf9 cell expression system by our labs) at 1 μg/ml. The plate was then blocked with 5% skimmed milk in PBS for 1 hr at 37°C, washed with PBS-T, loaded with 1/100 diluted mouse serum (0.06 ml/well), and incubated at room temperature for 2 hr. The plate was washed with PBS-T, treated with peroxidase-conjugated goat anti-mouse IgG antibody (1:10000; Abcam, UK), IgG1 (1:1000; Invitrogen), or IgG2a (1:1000; Invitrogen) for 1 hr at 37°C, washed, and loaded with TMB solution (Bio-Rad Laboratories, USA). Absorbance at 450 nm was measured using Epoch microplate reader (BioTek Instruments, USA). Each experiment was repeated three times.

### SARS-CoV2 neutralization assay

The SARS-CoV2 neutralizing assay was performed using SARS-CoV2 inside the Biosafety Level 3 facility (BSL3). Two weeks after immunizing the mice, serum samples were harvested, serially diluted 2-fold, and mixed 1:1 with SARS-CoV2 virus (prototype, Delta variant, or Omicron variant = 50TCID_50_). After a 1 hr incubation at 37°C, each virus-antibody mixture was added to Vero E6 cells, and the cells were incubated at 37°C for 3 days. The cells were then fixed and subjected to crystal violet staining. We used sera of patients who recovered after SARS-CoV2 infection as a positive control. The neutralizing assay was performed as described previously (9). Each experiment was repeated twice.

### SARS-CoV2 challenge in B6.cg-Tg (K18-ACE2) 2Prlmn/J mice

The challenge test was conducted 3 weeks after the last immunization. SARS-CoV2 was infected through the intranasal route (SARS-CoV2: 1×10^4^ TCID_50_/dose, SARS-CoV2 Delta variant: 1×10^4^ TCID_50_/dose, and SARS-CoV2 Omicron variant: 1×10^4^ TCID_50_/dose). Body weights were monitored after the challenge. The mice were sacrificed 14 days post-infection, and their lungs were harvested for histological staining and qPCR.

### Hematoxylin and eosin (H&E) staining

Mice were sacrificed, and lungs were harvested for Hematoxylin and eosin (H&E) staining. The lung tissue was fixed using 10% formalin, placed in paraffin, and sliced at 5-μm thickness. The lung tissue was deparaffinized according to our experimental protocol (9), stained with HE, and examined under an optical microscope. Each slides per animal was assessed to quantify the histological effects of the vaccine.

### Immunohistochemistry (IHC)

IHC was used to detect the expression of the N protein of SARS-CoV2. Prepared slides were deparaffinized in xylene and rehydrated through a graded ethanol series to distilled water. For SARS-CoV2-N, heat-induced epitope retrieval was performed by heating the slides in a pressure cooker on the steam setting for 10 min in 10 mM sodium citrate buffer and subsequently blocking the endogenous peroxidase using 3% hydrogen peroxide for 10 min. The slides were incubated with primary rabbit anti-SARS nucleocapsid protein antibody(NB100-56576, Novus, USA) overnight at 4°C and then with goat anti-rabbit IgG HRP (Abcam, UK) for 1 hr. Thereafter, 3,3’-diaminobenzidine (DAB) substrate (Takara, Japan) was added. All slides were counterstained with hematoxylin according to our experimental protocol (9) and dehydrated through a graded ethanol series to xylene. The slides were covered with mounting solution, and the HRP-oxidized brown precipitate of DAB was observed under an optical microscope.

### Quantitative real-time polymerase chain reaction (qRT-PCR)

SARS-CoV2 mRNA expression levels in lung tissue samples were analysed using qRT-PCR. Primers for the gene encoding N protein were used. RNA was extracted from each sample using the TRIzol^™^ reagent (Invitrogen) and used to synthesize cDNA with SuperScript^®^ Π reverse transcriptase (Invitrogen). qRT-PCR was performed using the SYBRGreen included in the BacPAK^™^ qPCR titration kit (Clontech, USA) and a StepOnePlus^™^ Real-Time PCR System (Applied Biosystems, USA). The total mRNA copy number was quantified using a standard curve prepared with serial dilutions of the pGEM-T vector containing a partial nucleocapsid gene of SARS-CoV2.

### IFN-γ ELISPOT assay

We assessed immunization and cellular immunity for various AcHERV-COVID19 vaccines in C57BL/6 or K18-hACE2 mice. Each group was immunized twice at a 4-week interval with the AcHERV-COVID19 vaccines. The production of interferon (IFN-γ) from splenocytes of immunized mice was detected by enzyme-linked immune spot (ELISPOT; BD Bioscience, USA) assay. Splenocytes were stimulated with inactivated SARS-CoV2 strains (prototype or Delta or Omicron; 7×10^5^ TCID50/well;). The color was developed using an AEC substrate reagent (BD Biosciences), and the spots were counted using an ELISPOT reader (AID ELISPOT Reader ver. 4; Germany).

### Cytokine mRNA measurements

RNA was extracted from spleen homogenates using the TRIzol^™^ reagent (Invitrogen), and cDNA was synthesized with SuperScript^®^ ⊓ reverse transcriptase (Invitrogen) according to the manufacturer’s protocol. Cytokine expression was determined with the SYBRGreen included in the Green Advantage qPCR Premix (Takara, Japan), using primers specific for TNF-a, IL-2, IL-4, and IL-10. The expression levels of the selected cytokine genes were normalized to that of GAPDH and converted to the relative expression the ratio (fold of induction), according to the following formula:

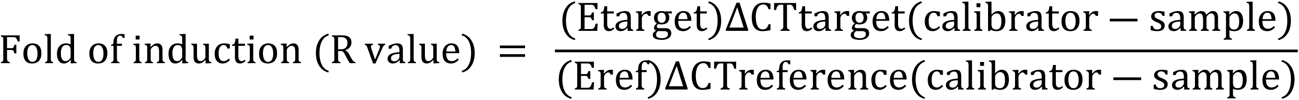

### Statistical analysis

All statistical analyses were performed using GraphPad Prism 8.0.2 (GraphPad

Software, Inc.), and data are presented as means ± standard deviation (SD). Comparison of data between groups was performed using an unpaired two-tailed t-test or one-way analysis of variance (ANOVA) followed by the Tukey–Kramer post-hoc test. *P*-values <0.05 were considered statistically significant.

## Acknowledgments

The pathogenic resources (NCCP No.43326, NCCP No.43390 NCCP No.43408, SARS-CoV2 strains) for this study were provided by the National Culture Collection for Pathogens. This research was supported by a grant from the Korea Health Technology R&D Project through the Korea Health Industry Development Institute (KHIDI), funded by the Ministry of Health & Welfare, Republic of Korea (grant No.HQ21C0264, HV22C0263) and supported by a grant (22183MFDS443) from Ministry of Food and Drug Safety in 2022.

## Author Contribution

Y Jang, H Cho: Construction of the AcHERVs, Data curation, JM Chun, KH Park, A Nowakowska: In vivo analysis (ABL3, BL3), JH Kim: ELISA assay, HD Lee, CY Lee: Histological assay, HJ Lee, Y Han, HY Shin: material supply and data analysis, YB Kim: Corresponding author, Design, and writing.

## Supporting information

**S1 Fig. Cellular immune response in AcHERV-COVID19S vaccinated mouse splenocytes** (A) Induction of SARS-CoV2-specific T cells. The splenocytes were harvested 2 weeks after the last immunization and stimulated with inactivated SARS-CoV2 virus for 24 hr. The number of IFN-γ-producing SARS-CoV2 spike-specific CD8+ T cells was determined using an ELISPOT assay. (B) Quantification of T cell related cytokines by qPCR. The mRNA expression of cytokines (Th1: TNF-α, IL-2; Th2: IL-4) was determined by qRT-PCR, normalized to GAPDH, and compared to NTC controls in spleen homogenates 2 weeks after last immunization. Data were analyzed with unpaired two-tailed t-test or one-way analysis of variance (ANOVA) followed by the Tukey-Kramer post analysis tests. NS: P > 0.05, *: P < 0.05, **: P < 0.01, ***: P < 0.001, ****: P < 0.0001.

**S2 Fig. Schematic diagrams of the transfer plasmid (Ad-COVID19S/ COVID19D) for the construction of recombinant adenovirus and expression of 293T.** (A) The recombinant adenoviruses were constructed to contain SARS-CoV2 prototype or delta S genes under the CMV promoters. (B) The expression of SARS-CoV2 S proteins was analyzed by western blotting in 293 T cells. Specific rabbit anti-SARS-CoV2 S sera were used as a primary antibody. The molecular weight of SARS-CoV2 S full, and S1 are approximately 180kDa, and 100kDa respectively. (-): uninfected cells; Lane 1: Ad-COVID19S; Lane 2: Ad-COVID19D.

**S1 Table. Immunization and Challenge test for AcHERV-COVID19S in ACE2 mice.** 14 days after the last immunization, mice were challenged with SARS-CoV2 prototype through the intranasal route (1×10^4^ TCID_50_/dose). The mice were observed daily weight change for 13 days after challenge.

**S3 Table. Immunization and Challenge test for AcHERV-COVID19S and AcHERV-COVID19D in ACE2 mice.** 13 days after the last immunization, mice were challenged with SARS-CoV2 Delta through the intranasal route (1×10^4^ TCID50/dose). The mice were observed daily weight change for 13 days after challenge.

